# *Rbpj* deletion in hepatic progenitor cells attenuates endothelial responses and fibrosis in DDC-fed mice

**DOI:** 10.1101/2024.04.13.589277

**Authors:** Sanghoon Lee, Lu Ren, Weiwei Li, Aditi Paranjpe, Ping Zhou, Andrew Potter, Stacey S. Huppert, Soona Shin

## Abstract

**Background:** As the role of hepatic progenitor cells (HPCs) constituting ductular reactions in pathogenesis remains ambiguous, we aimed to establish the in vivo cause-and-effect relationship between HPCs and chronic liver disease progression. We previously demonstrated that peritumoral ductules are associated with angiogenesis in liver tumors and forkhead box L1 (Foxl1)-expressing murine HPCs secrete angiogenic factors in vitro. Therefore, we hypothesized that HPCs are capable of remodeling the portal vascular microenvironment and regulating overall liver disease progression, and this function of HPCs is dependent on recombination signal binding protein for immunoglobulin kappa J region (RBPJ), a key effector of the Notch signaling pathway.

**Methods and Results:** We generated HPC-specific *Rbpj* conditional knockout mice (CKO) using *Foxl1-Cre* and treated them with the DDC diet to induce chronic liver disease. CKO mice exhibited a significant reduction in serum levels of liver injury markers, ductular reactions, vascular and fibrotic areas, and hepatic expression of fibrosis and inflammation markers compared to control mice (WT). Single-nucleus RNA sequencing comparing CKO and WT livers detected transcriptome changes across multiple cell types, including endothelial cells, hepatic stellate cells, and cholangiocytes. Expression of several reactive cholangiocyte markers, including vascular cell adhesion molecule 1 (VCAM1), in HPCs was significantly downregulated in response to anti-*Rbpj* shRNAs in vitro. Immunofluorescence analysis indicated that the percentage of VCAM1+ cells was reduced in both HPC and cholangiocyte populations in CKO compared to WT in vivo.

**Conclusions:** Our findings reveal *Rbpj*-dependent expression of reactive cholangiocyte markers in HPCs and demonstrate that *Rbpj* deletion in HPCs attenuates not only endothelial responses but also liver injury, fibrosis, and VCAM1 expression in cholangiocytes, highlighting the crucial role of HPCs in pathogenic progression.

## INTRODUCTION

Hepatic progenitor cells (HPCs) are facultative stem cells that are mainly derived from normal cholangiocytes activated in response to liver injury.[1] HPCs form ductular reactions and peritumoral ductules detected in a variety of chronic liver diseases and cancer.[2–6] However, their causal role in disease progression has yet to be established. Previous studies reported that HPCs express paracrine factors involved in angiogenesis, fibrosis, and inflammation in the context of various liver diseases such as primary biliary cirrhosis and pediatric metabolic dysfunction-associated steatotic liver disease.[7,8] We also reported that HPCs in peritumoral ductules in the livers of patients with liver cancer express angiogenic factors, and the number and proliferation of HPCs, as well as the extent of angiogenic factor expression by HPCs, correlate with intra- and peritumoral angiogenesis and intratumoral proliferation.[4]

Forkhead box L1 (Foxl1)-expressing HPCs and their descendants can be labeled with yellow fluorescent protein (YFP) using the *Foxl1-Cre* transgenic mouse in multiple models of liver disease[9–11] including 3,5-diethoxycarbonyl-1,4-dihydrocollidine (DDC)-induced chronic liver disease.[12] HPCs isolated from DDC-fed *Foxl1-Cre;Rosa^YFP^* mice secrete numerous paracrine factors involved in angiogenesis, fibrosis, and tumorigenesis.[13] Our in vitro data also indicated that HPCs communicate with endothelial cells in a paracrine manner to stimulate endothelial proliferation, tubulogenesis, and gene expression, implying their potential roles in angiogenesis.[13] Several studies have reported an association of angiogenesis with other pathogenic mechanisms such as fibrosis and inflammation in the context of various liver diseases, including hepatocellular carcinoma, metabolic dysfunction-associated steatotic liver disease, and cholangiopathies.[14,15] Association between HPCs and angiogenesis has also been reported, but these studies were based on correlative data or global gene disruption affecting multiple cell types.[4,7,16] Therefore, the causal role of HPCs in altering the endothelium of diseased liver using specific modulation of HPCs remains to be tested. Thus, the current study aims to establish the cause-and-effect relationship between HPCs and DDC-induced endothelial responses using conditional gene modulation specific for HPCs. DDC mainly induces biliary injury but also causes periportal hepatocytic injury, and it is a widely used model for inducing ductular reactions and simulating cholangitis and biliary fibrosis.[12,17,18]

It has been demonstrated that recombination signal binding protein for immunoglobulin kappa J region (RBPJ), the effector molecule of the Notch signaling pathway, is an important regulator of biliary morphogenesis.[19] However, the study used the albumin promotor as the driver, which is active in hepatocytes, biliary cells, as well as the fetal precursor hepatoblasts. Therefore, the specific requirement of RBPJ for the proliferation and function of Foxl1-expressing postnatal HPCs during pathogenic progression remains largely unclear. As such, we aim to address how *Rbpj* gene deletion in HPCs affects communication of HPCs with other hepatic cells. While our previous study focused on HPC expression of angiogenic factors,[13] (1) based on the close association between vascular responses, fibrosis, and ductular reactions in diseased livers, (2) the involvement of reactive cholangiocytes in chronic liver disease progression, and (3) our previous observation that the transcriptome of HPCs clusters more closely with cholangiocytes compared to hepatocytes,[9] we hypothesize that HPCs retain certain cholangiocyte characteristics despite their bipotential differentiation capability and promote not only angiogenesis but also overall progression of liver disease by expressing markers of reactive cholangiocytes. We generated conditional knockout mice in which *Rbpj* is specifically deleted in HPCs by crossing *Foxl1-Cre;Rosa^YFP^* mice[9] with *Rbpj^F/F^*mice.[20] This allowed us to determine the impact of HPC modulation on the liver microenvironment and the requirement of RBPJ in this process.

## MATERIALS AND METHODS

### Animal experiments

The sources of *Foxl1-Cre*, *Rosa^YFP^*, and *Rbpj^F/F^* mice are as described previously.[20–22] *Foxl1-Cre;Rosa^YFP^;Rbpj^F/+^* mice on C57BL/6J background were used for breeding to generate *Foxl1-Cre;Rosa^YFP^;Rbpj^F/F^* conditional knockout mice (CKO) and *Foxl1-Cre;Rosa^YFP^;Rbpj^+/+^* wild-type littermate control mice (WT). The 10-to 11-week-old male mice were fed a diet supplemented with 0.1% DDC (Envigo) for 4 weeks. All procedures were approved by the Institutional Animal Care and Use Committee at Cincinnati Children’s Hospital Medical Center.

### Histological analysis of liver sections

Formalin-fixed paraffin-embedded tissue sections were subjected to immunofluorescence staining and quantification as described previously.[10] Six to eight random images centered on the portal area were captured for each mouse. HPCs were defined as *Foxl1-Cre*/YFP-labeled cells within the cytokeratin 19 (CK19)-positive biliary epithelial population.[9,10] The following primary antibodies were used: GFP (Aves Labs GFP-1020), RBPJ (Cosmo Bio USA SIM-2ZRBP2), CK19 (Abcam ab52625), Ki67 (Thermo Fisher Scientific 14-5698-82), CD31 (Abcam ab28364), CD34 (Abcam ab81289), αSMA (Abcam ab7817), GFAP (Thermo Fisher Scientific 14-9892-82), and VCAM1 (R&D Systems AF643). TSA Plus Cyanine 3 System (PerkinElmer NEL744001KT) was used to amplify signals generated using antibodies for RBPJ, CD31, CD34, αSMA, and GFAP. Nikon Eclipse Ti microscope was used for imaging. Nikon A1 inverted LUNV confocal microscope was used to determine the expression of RBPJ and VCAM1 by HPCs and cholangiocytes. Trichrome staining was performed using Trichrome Staining Kit (Roche Diagnostics 860–031). Whole liver sections were scanned with a Leica AT2 digital slide scanner (Leica Biosystems) and five random images were captured for each mouse. ImageJ was used for cell counting and quantification of marker-positive areas.[23]

### Cell culture

A clonally expanded HPC line that we previously established from a DDC-fed mouse was used.[9] HPC culture medium was prepared as described in our previous studies.[9,13] HPCs were treated with lentiviral vectors expressing the non-silencing control and each of the 2 different shRNAs targeting the mouse *Rbpj* gene (VectorBuilder). Cells were maintained in the culture medium containing 5 μg/mL puromycin to select transduced cells. Cells were grown to 70% confluency and harvested for subsequent analysis.

### Quantitative polymerase chain reaction

RNA isolation, cDNA synthesis, and quantitative polymerase chain reaction (qPCR) were performed, and data were normalized to *Tbp* as described previously.[13] Primer sequences are available in Supplemental Table S1.

### Serum biochemistry

Blood drawn from the inferior vena cava was incubated at room temperature for 30 minutes and placed on ice for 1 hour. Serum was collected by centrifugation at 2,000g for 10 minutes at 4°C and analyzed using Mouse AST ELISA Kit (Abcam ab263882) and Mouse ALP ELISA Kit (Abcam ab285274).

### Statistical analysis

Student’s t-tests or analysis of variance followed by multiple comparison tests were performed where applicable to detect the significance of differences between groups (GraphPad Prism 10). A p-value of <0.05 was considered statistically significant.

Please refer to the Supplemental Materials and Methods for additional information.

## RESULTS

### Ductular reactions are significantly reduced in mutant mice

To establish the cause-and-effect relationship between HPCs and vascular responses in cholestatic liver disease, we aimed to test the hypothesis that HPC-specific disruption of the *Rbpj* gene attenuates ductular reactions. It has been reported that systemic inhibition of Notch signaling using a pharmacologic inhibitor or *Alb-Cre*-mediated deletion of *Rbpj* in all hepatic epithelial populations reduced the number of HPCs and impaired biliary tubulogenesis in DDC-fed mice.[19,24] However, whether conditional *Rbpj* knockout using a promoter specific for HPCs impacts ductular reactions remains to be determined. We previously demonstrated that Foxl1 is a specific marker for facultative HPCs derived from normal cholangiocytes activated in response to various types of liver injury.[5,6,9] Since Foxl1 is not detected in uninjured healthy livers and is only expressed in HPCs upon injury, the *Foxl1-Cre* transgenic mouse offers a valuable way to modulate gene expression of HPCs without affecting normal cholangiocytes. Therefore, to determine the impact of *Rbpj* knockout on ductular reactions and HPC number, *Foxl1-Cre;Rosa^YFP^;Rbpj^F/F^* (CKO) mice and *Foxl1-Cre;Rosa^YFP^;Rbpj^+/+^*(WT) control mice were treated with the DDC diet for 4 weeks (Figure 1A). We performed immunostaining analysis of liver sections and quantified the percentage of RBPJ-expressing cells within the cytokeratin 19 (CK19)-expressing biliary population, a subset of which are *Foxl1-Cre*/YFP-labeled HPCs. While the extent of RBPJ expression was not different between CK19+YFP+ HPCs and CK19+YFP-cholangiocytes in DDC-fed WT mice (WT-DDC), CK19+YFP+ HPCs in DDC-fed CKO mice (CKO-DDC) displayed a reduced percentage of RBPJ-expressing cells compared to CK19+YFP-cholangiocytes in CKO-DDC as well as CK19+YFP+ HPCs in WT-DDC, confirming that RBPJ is specifically deleted in HPCs of CKO mice (Figures 1B, C).

**Figure 1.**
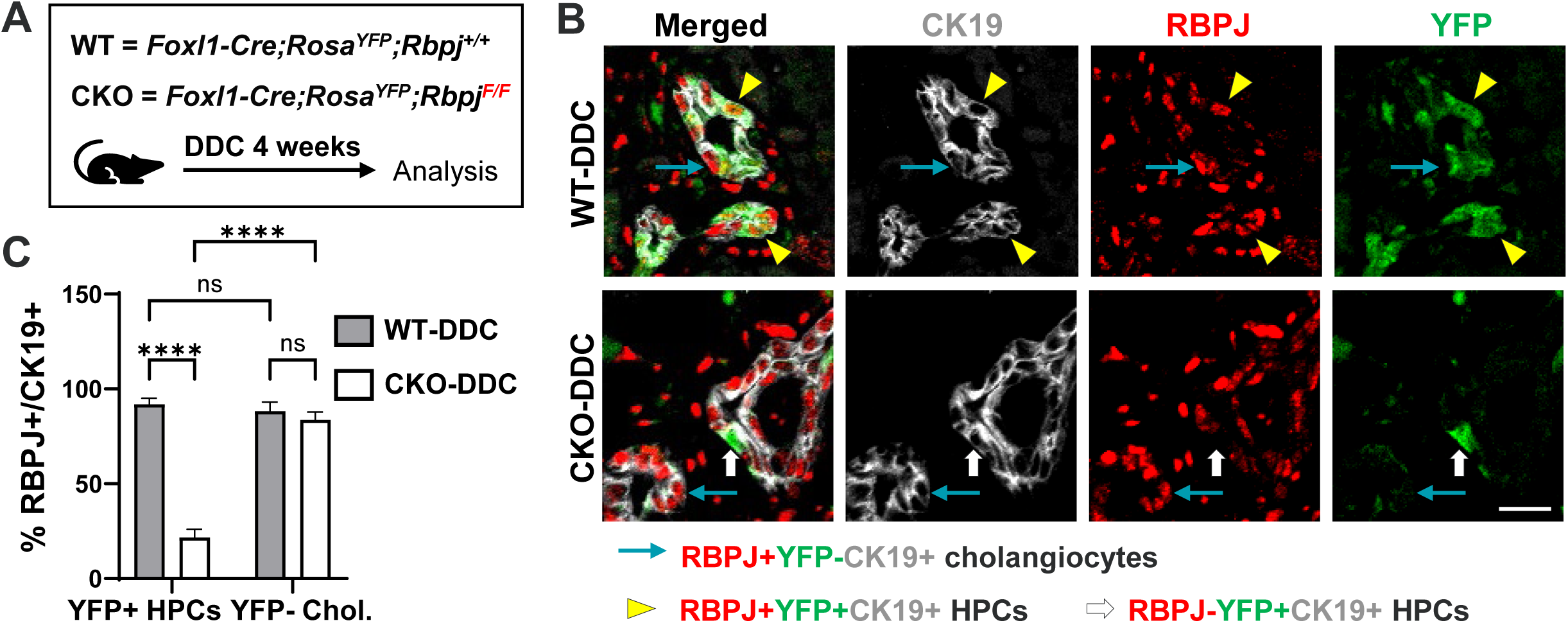
Conditional deletion of *Rbpj* in HPCs using *Foxl1-Cre*. (A) Schematic representation of the experimental strategy. 10-to 11-week-old male mice were treated with the DDC diet for 4 weeks. (B) Confocal immunofluorescence analysis. Top panels: RBPJ protein is expressed in YFP+CK19+ HPCs (yellow arrowheads), YFP-CK19+ cholangiocytes (blue arrows), and surrounding mesenchymal cells in portal areas of WT livers. Bottom panels: RBPJ protein expression is lost specifically in HPCs (white arrows) of CKO livers. WT-DDC and CKO-DDC refer to DDC-fed WT and CKO mice, respectively. Scale bar, 20 μm. (C) Quantification of the percentage of RBPJ-expressing cells within the CK19+ biliary population in portal areas. Chol. = cholangiocytes. Data are expressed as mean ± SE; n = 5 mice per group; **** p < 0.0001; ns, not significant. Abbreviations: CK19, cytokeratin 19; CKO, conditional knockout; DDC, 3,5-diethoxycarbonyl-1,4-dihydrocollidine; HPC, hepatic progenitor cell; RBPJ: recombination signal binding protein for immunoglobulin kappa J region; WT, wild-type; YFP, yellow fluorescent protein.

To determine the effect of HPC-specific *Rbpj* deletion on ductular reactions, we quantified the number of CK19-expressing cells in the portal areas. Treatment of WT mice with DDC led to a dramatic induction of the total number of CK19+ cells per area compared to age-matched chow-fed mice (WT-Chow), while it was significantly reduced in CKO-DDC compared to WT-DDC, indicative of attenuated ductular reactions in response to conditional *Rbpj* deletion (Figure 2A). Since the total number of CK19+ cells in DDC-fed mice is the sum of YFP+ HPCs and YFP-cholangiocytes, we quantified the number of these two cell populations per area in DDC-fed mice (Figure 2B) to determine the cell type contributing to attenuated ductular reactions. Both CK19+YFP+ HPCs and CK19+YFP-cholangiocytes displayed a trend toward decreased numbers in CKO-DDC compared to WT-DDC. While changes in each population did not reach statistical significance, the sum of them did, leading to reduced ductular reactions in CKO mice. Quantification of Ki67+ cells indicated that the percentages of proliferating CK19+YFP+ HPCs and CK19+YFP-cholangiocytes are reduced in CKO-DDC compared to WT-DDC (Figure 2C) in line with the reduced cell numbers, although the difference did not reach significance in the cholangiocyte population.

**Figure 2.**
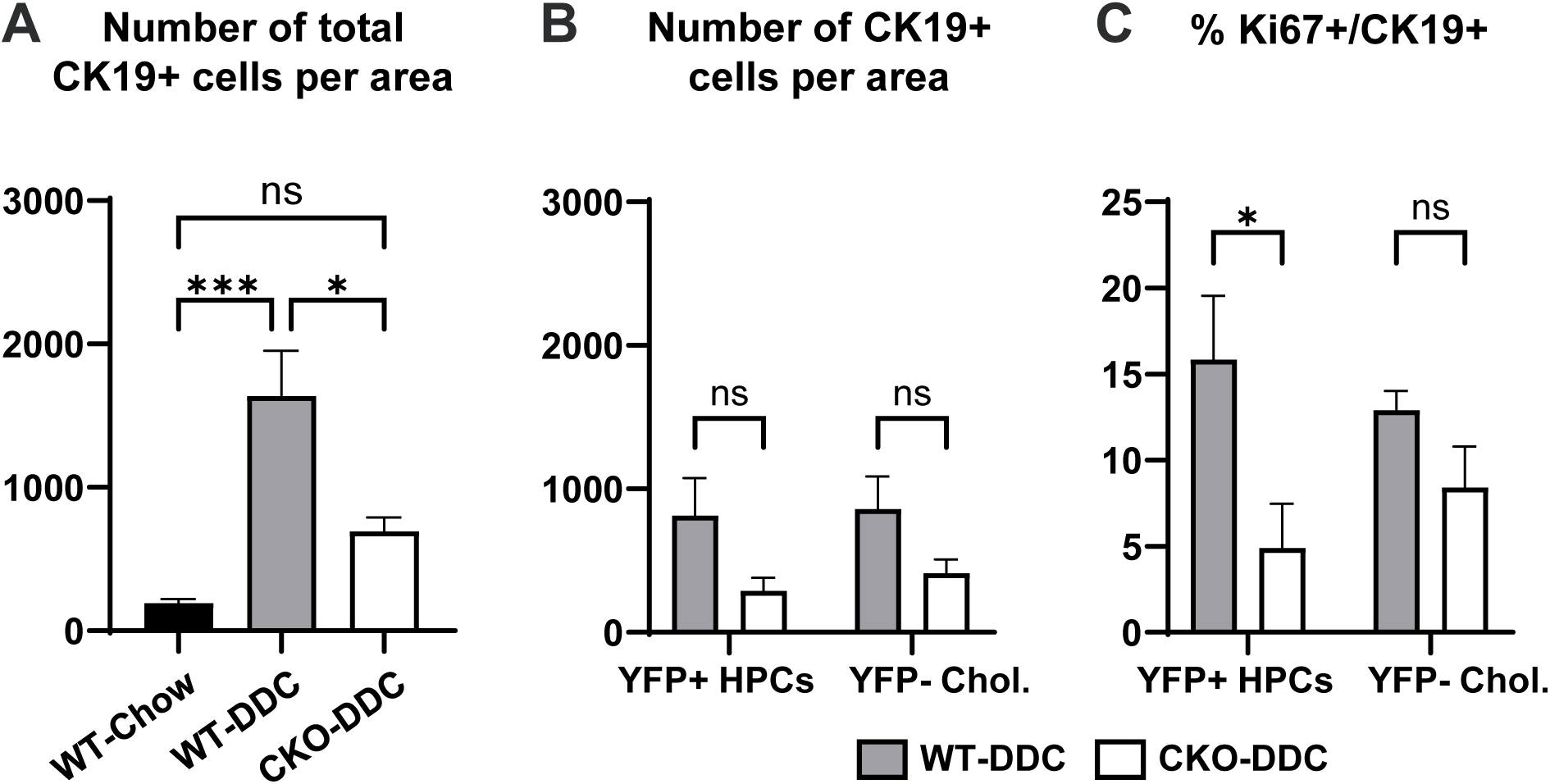
Ductular reactions are attenuated in CKO mice. (A) Number of CK19+ cells per portal area in the livers of chow-fed WT mice (WT-Chow), WT-DDC, and CKO-DDC. (B) Number of CK19+YFP+ HPCs and CK19+YFP-cholangiocytes (Chol.) per portal area in WT-DDC and CKO-DDC livers. (C) Percentage of Ki67+ cells in relation to the total number of CK19+YFP+ HPCs and CK19+YFP-cholangiocytes in the portal areas of WT-DDC and CKO-DDC livers. All data are expressed as mean ± SE; n = 4-5 mice per group; * p < 0.05; *** p < 0.001; ns, not significant. Abbreviations: CKO, conditional knockout; CK19, cytokeratin 19; DDC, 3,5-diethoxycarbonyl-1,4-dihydrocollidine; HPC, hepatic progenitor cell; WT, wild-type.

### Serum levels of liver injury markers are decreased in CKO compared to WT

To determine how conditional *Rbpj* knockout impacts liver injury, we analyzed the percent body weight loss in response to 4 weeks of DDC treatment and the liver weight/body weight ratio (Figure 3A). Neither the extent of body weight loss nor the liver weight/body weight ratio reached statistically significant differences between WT-DDC and CKO-DDC, while the liver weight/body weight ratio was increased in WT-DDC compared to WT-Chow in line with data reported in the literature.[25,26] The serum levels of liver injury markers were also assessed (Figure 3B). Previous studies demonstrated that the DDC diet induces not only cholestatic injury but also periportal hepatocytic injury.[12,18] Indeed, the serum levels of both the hepatocytic injury marker aspartate aminotransferase (AST) and the cholestatic injury marker alkaline phosphatase (ALP) were significantly increased in WT-DDC compared to WT-Chow, while there was a statistically significant decrease in AST and ALP levels in CKO-DDC compared to WT-DDC. These results indicate that HPC-specific *Rbpj* deletion attenuates DDC-induced liver injury.

**Figure 3.**
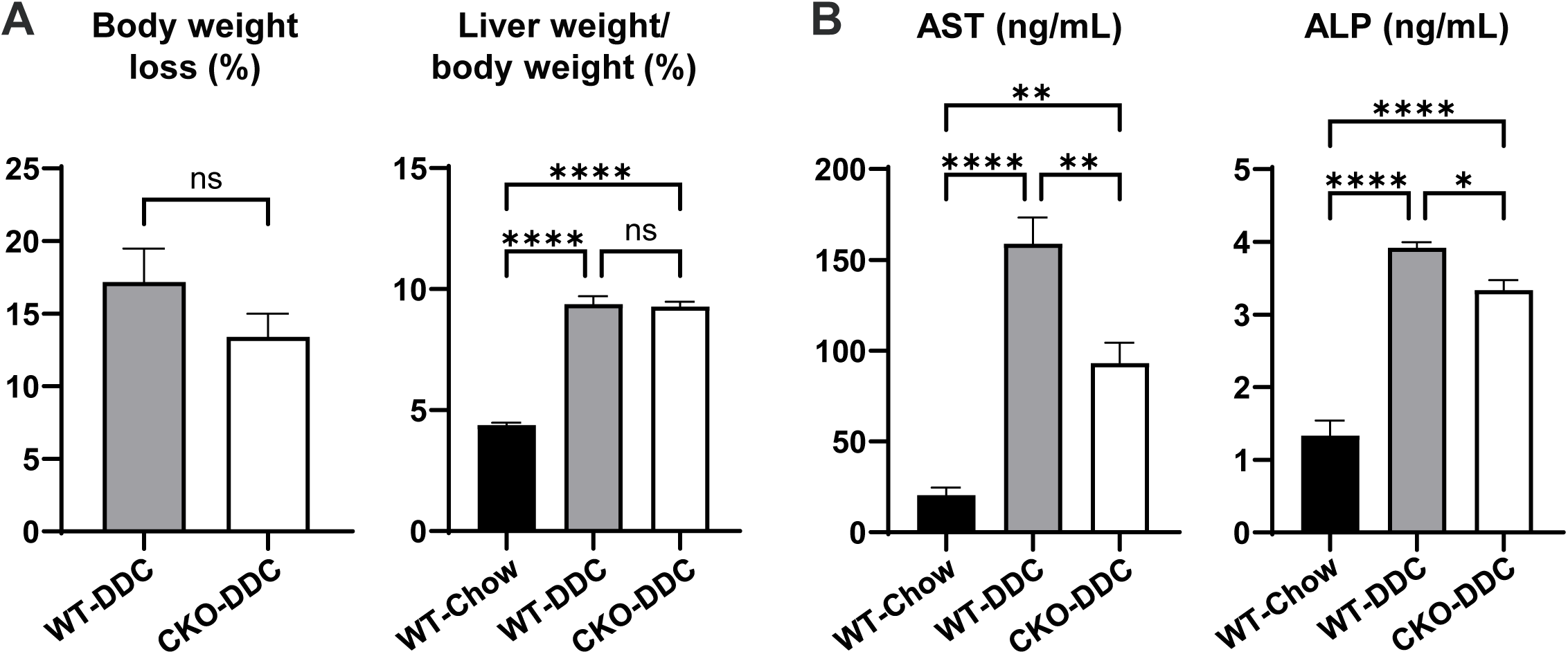
Serum levels of liver injury markers. (A) The body weight loss (relative to initial body weight) after 4 weeks of DDC diet treatment and the liver weight/body weight ratio are shown. Data are expressed as mean ± SE; WT-Chow (n = 5); WT-DDC (n = 11); CKO-DDC (n = 11). (B) Serum biochemistry. Data are expressed as mean ± SE; WT-Chow (n = 6); WT-DDC (n = 9); CKO-DDC (n = 9). * p < 0.05; ** p < 0.01; **** p < 0.0001; ns, not significant. Abbreviations: ALP, alkaline phosphatase; AST, aspartate aminotransferase; CKO, conditional knockout; DDC, 3,5-diethoxycarbonyl-1,4-dihydrocollidine; WT, wild-type.

### Rbpj deletion in HPCs attenuates portal microenvironment alterations, fibrosis, and inflammation

Since the DDC diet predominantly affects portal areas by inducing biliary fibrosis and ductular reactions,[12,18] we assumed that analysis of histopathological markers focusing on the local microenvironment would reveal additional insights into how HPCs are involved in disease progression. Therefore, we performed immunostaining analysis of portal areas in which HPCs are detected. Based on our previous study suggesting that HPCs communicate with the endothelial compartment[4,13], we investigated the impact of conditional *Rbpj* deletion on areas occupied by endothelial cells (Figure 4A, B). CD31 is a pan-endothelial marker.[27] CD34 is expressed in vascular endothelial cells but not in liver sinusoidal endothelial cells (LSECs) in healthy livers; however, its expression is detected in LSECs in diseased livers. CD34 is widely used to measure the microvessel density in liver disease[28] and is also expressed by a subset of fibroblasts and fibrocytes.[29] Our data indicate that the areas positive for CD31 and CD34 are dramatically reduced in CKO-DDC compared to WT-DDC, suggesting that the portal microenvironment responds to conditional *Rbpj* modulation specific for HPCs.

**Figure 4.**
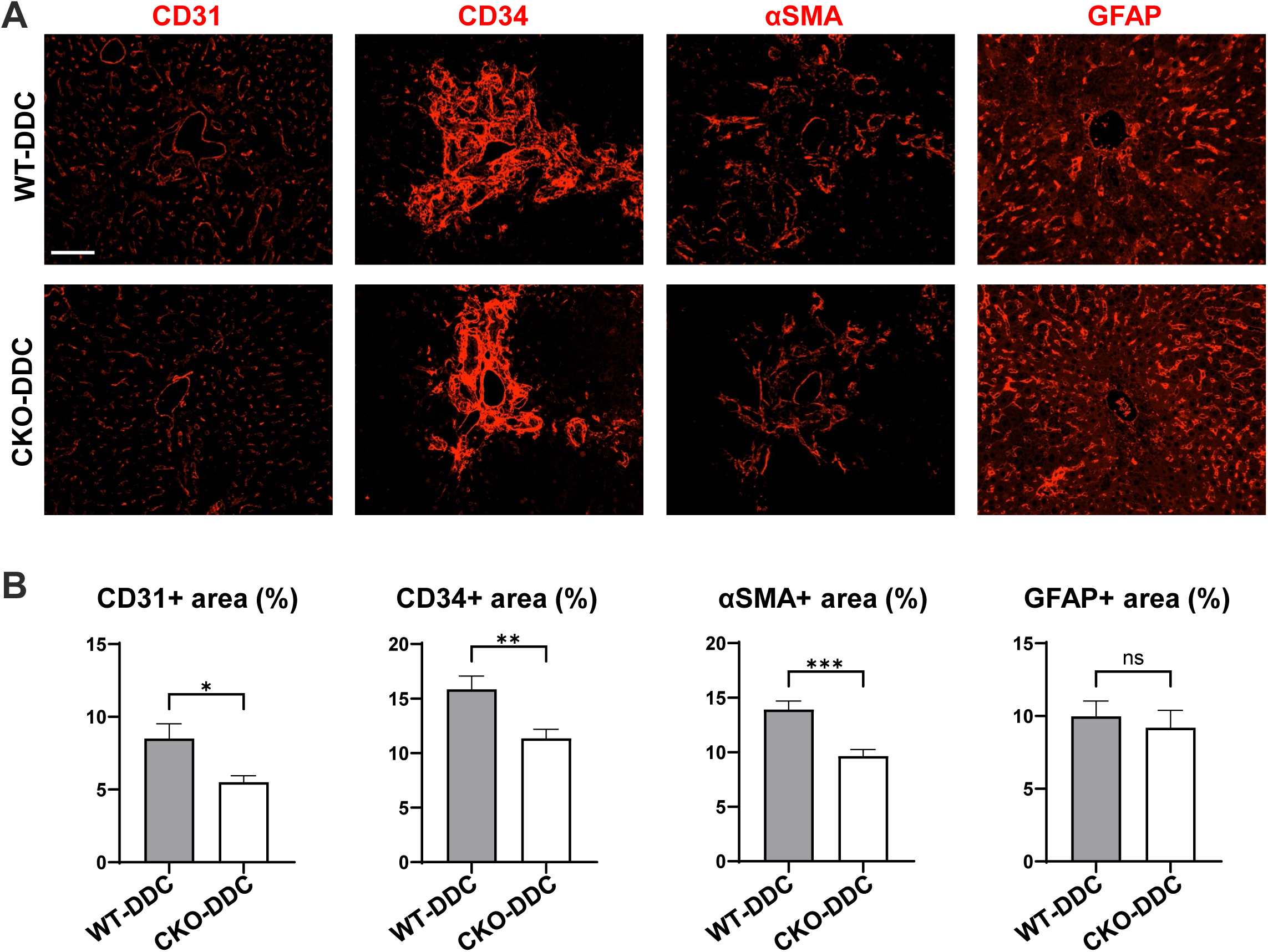
Portal endothelial and fibrotic areas are reduced in CKO-DDC compared to WT-DDC. (A) Immunofluorescence analysis of portal areas. Scale bar, 100 μm. (B) Quantification of marker-positive portal areas. All data are expressed as mean ± SE; n = 11 mice per group. * p < 0.05; ** p < 0.01; *** p < 0.001; ns, not significant. Abbreviations: αSMA, alpha smooth muscle actin; CKO, conditional knockout; DDC, 3,5-diethoxycarbonyl-1,4-dihydrocollidine; GFAP, glial fibrillary acidic protein; WT, wild-type.

Based on the reported association between angiogenesis and other pathogenic mechanisms such as liver fibrosis and inflammation,[14,15] and reduced levels of serum injury markers in CKO mice (Figure 3B), we hypothesized that altered endothelial responses in CKO mice are attributable to attenuated liver disease progression in CKO-DDC compared to WT-DDC. To this end, we performed immunofluorescence analysis of portal areas for markers of fibrosis. Alpha smooth muscle actin (αSMA) is a well-established marker of myofibroblasts. Glial fibrillary acidic protein (GFAP) is expressed in quiescent hepatic stellate cells (HSCs).[30,31] Our data indicate that the area positive for αSMA is substantially reduced in CKO-DDC compared to WT-DDC, while the GFAP-positive area remained unaffected (Figures 4A, B). These results suggest that modulation of HPCs does not affect the total number of quiescent HSCs but rather inhibits the activation of precursor cells (HSCs or portal fibroblasts) into myofibroblasts.

Since our immunofluorescence analysis focused on the portal areas, we also attempted to capture the extent of fibrosis and inflammation of the whole liver. Indeed, trichrome analysis, in which images were randomly captured regardless of liver zones, indicated that the expansion of the fibrosis area in WT-DDC compared to WT-Chow is attenuated in CKO-DDC (Figures 5A, B). qPCR analysis of mouse liver tissues indicated that the mRNA levels of fibrosis markers (Figure 5C) and inflammation markers (Figure 5D) are upregulated in WT-DDC compared to WT-Chow and downregulated in CKO-DDC compared to WT-DDC. In summary, our results in Figures 2-5 establish the causal relationship between HPCs and not only portal vascular responses but also overall liver disease progression in DDC-fed mice.

**Figure 5.**
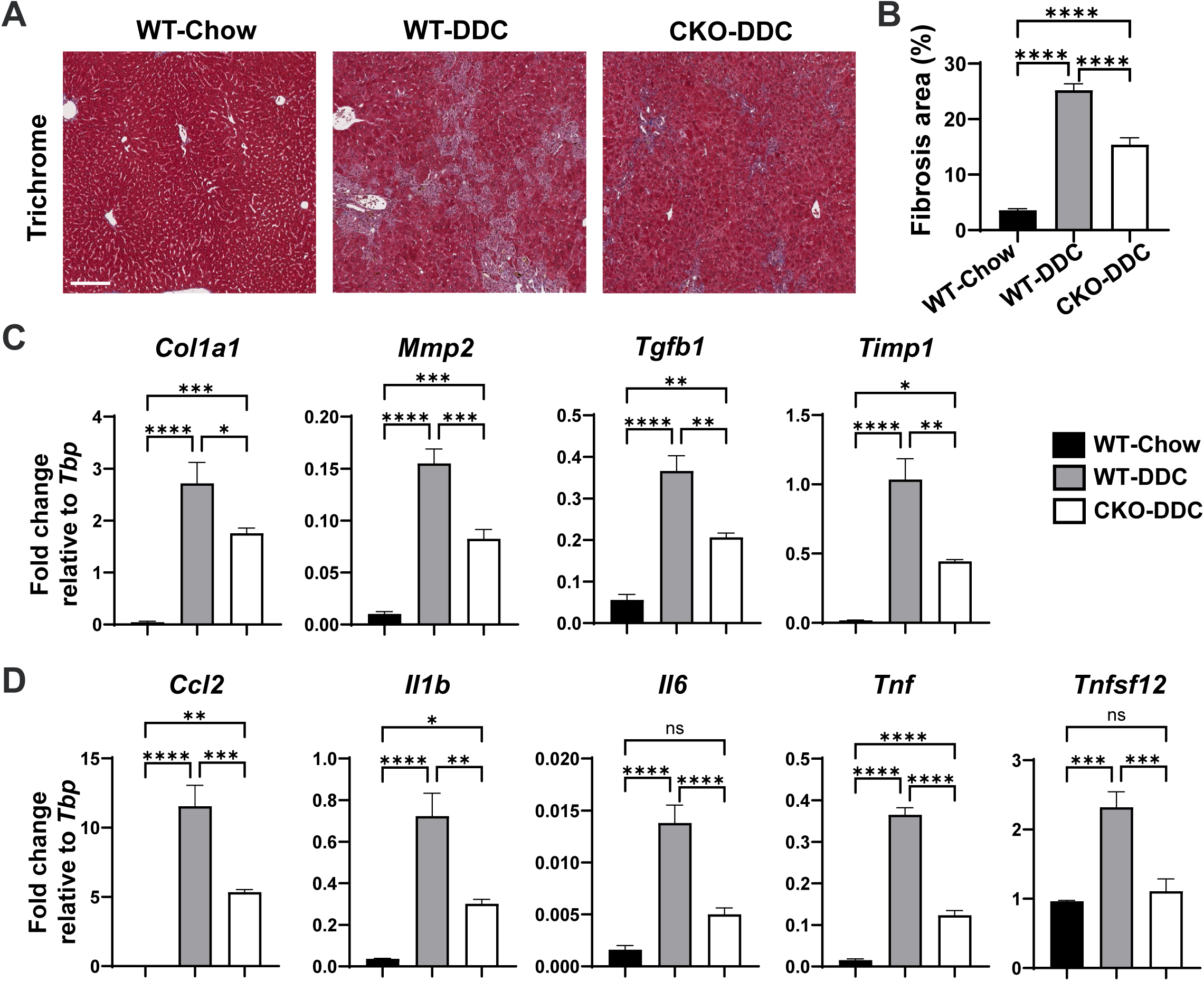
Markers of liver fibrosis and inflammation are downregulated in CKO-DDC compared to WT-DDC. (A) Trichrome staining of liver sections. Scale bar, 150 μm. (B) Quantification of Trichrome Blue-stained collagen area. (C, D) qPCR analysis of mouse livers. The mRNA levels of markers of fibrosis (C) and inflammation (D) are shown. All data are expressed as mean ± SE; n = 5 mice per group. * p < 0.05; ** p < 0.01; *** p < 0.001; **** p < 0.0001; ns, not significant. Abbreviations: CKO, conditional knockout; DDC, 3,5-diethoxycarbonyl-1,4-dihydrocollidine; qPCR, quantitative polymerase chain reaction; WT, wild-type.

### Single-nucleus RNA sequencing indicates altered gene expression in multiple cell clusters upon conditional modulation of HPCs

To determine whether the transcriptome changes in the mouse liver align with our immunostaining analysis indicating that conditional deletion of *Rbpj* in HPCs affects other cell types, we performed single-nucleus RNA sequencing (sNuc-seq) of the livers of WT-DDC (n = 2) and CKO-DDC (n = 2). A total of 6 cell clusters were identified in both WT and CKO mice (Supplemental Figures S1A-C). We identified cluster-specific markers in an unbiased manner by determining highly expressed and unique genes in one cluster versus all other clusters using Seurat (Supplemental Materials and Methods, Supplemental Table S2). Annotation of each cluster was performed by determining the overlaps between our markers in Supplemental Table S2 and those reported by Liang and colleagues who performed comprehensive single-cell transcriptomic analysis of mouse livers (Supplemental Table S3).[32] Two hepatocyte clusters were identified, Cluster 0 (Hepatocytes-1) and Cluster 2 (Hepatocytes-2). Gene set enrichment analysis (GSEA) of cluster-enriched marker genes (Supplemental Table S3) indicated that each of the hepatocyte clusters is segregated based on distinct metabolic functions (Supplemental Table S4). Non-parenchymal cell (NPC) populations detected include: Cluster 1 - myeloid cells, Cluster 3 - cholangiocytes, Cluster 4 - endothelial cells, Cluster 5 - HSCs (Supplemental Figures S1A-C). However, a cluster specific for *Foxl1*-expressing HPCs was not detected in our sequencing data.

While analysis of differentially expressed genes between WT-DDC and CKO-DDC indicated that conditional modulation of HPCs impacts gene expression in many clusters (Supplemental Table S5), we were particularly interested in GSEA results (Supplemental Tables S6-S11) from genes downregulated in the CKO-DDC cells contributing to each cluster, since they can be inferred as processes activated when WT HPCs are present. Cluster 4 expressed vascular endothelial cell markers such as *Ptprb* and the LSEC marker *Stab2* (Supplemental Table S3, Supplemental Figure S1B).[33] GSEA of genes downregulated in CKO-DDC compared to WT-DDC in Cluster 4 identified gene sets involved in endothelial proliferation and survival, including vascular endothelial growth factor signaling, DNA replication, and hedgehog signaling (Supplemental Figure S1D, Supplemental Table S10).[34,35] In line with attenuated ductular reactions and fibrosis upon HPC-specific *Rbpj* deletion (Figures 2, 4, 5), biological processes related to proliferation and functions of cholangiocytes and HSCs were downregulated in CKO-DDC compared to WT-DDC (Supplemental Figure S1D, Supplemental Tables S9, S11). In summary, our data indicate that conditional genetic modulation of HPCs induces transcriptomic alterations across multiple cell types.

### HPCs express markers of reactive cholangiocytes

Because our sNuc-seq did not detect a separate *Foxl1*-expressing HPC population, to understand the mechanism by which HPC-specific *Rbpj* deletion impacts other hepatic cell types and overall disease progression, we revisited our previously published microarray data GSE28892.[9] According to this previous study, comparison of HPCs from DDC-fed mice to cholangiocytes from chow-fed mice and hepatocytes from chow-fed mice indicated that HPCs express markers of both cholangiocytes and hepatocytes, and the transcriptome of HPCs clusters closer to cholangiocytes compared to hepatocytes, suggesting that HPCs retain certain cholangiocyte characteristics despite their bipotential nature. Based on the literature indicating that cholangiocytes in injured liver express secretory factors involved in pathogenic progression[36–38] and our observation that HPCs secrete angiogenic factors in vitro,[13] we hypothesized that HPCs express markers of reactive cholangiocytes. Our data in Supplemental Figure S2 are based on two different sets of microarray analysis. Supplemental Figure S2A compares HPCs from WT-DDC to normal cholangiocytes and hepatocytes isolated from WT-Chow. Supplemental Figure S2B compares HPCs from WT-DDC to NPCs from the same DDC-fed mice, the latter consisting of the remaining cells after exclusion of both HPCs and hepatocytes. These data indicate that HPCs express high levels of many reactive cholangiocyte markers[36–38] in at least one comparison.

### Reactive cholangiocyte marker expression in HPCs is dependent on Rbpj

To understand the mechanism underlying reduced endothelial and fibrotic areas in CKO-DDC compared to WT-DDC, we investigated whether knocking down *Rbpj* in HPCs affects expression of reactive cholangiocyte markers[36–38] in vitro. A clonally expanded HPC line previously established from a DDC-fed mouse[9] was used to circumvent the effect of cell heterogeneity. HPCs were treated with lentiviral vectors expressing two different anti-*Rbpj* shRNAs (shRNA-1 and shRNA-2) as well as a non-silencing control and subjected to puromycin selection. qPCR analysis indicated that both shRNAs downregulate the mRNA level of *Rbpj* as well as *Cdon, Notch4*, and *Sox2* which are reported to be downstream of Notch signaling and direct targets of RBPJ (Figure 6A).[39,40] While the expression level of *Ccl2* and *Vegfa* remained unchanged or even slightly increased in response to *Rbpj* knockdown, many markers including *Angpt2*, *Il6*, *Slit2*, *Tnf*, and *Vcam1* known to promote ductular reactions, angiogenesis, fibrosis, and inflammation were significantly downregulated in shRNA-1- and shRNA-2-treated cells compared to the control group (Figure 6B).

**Figure 6.**
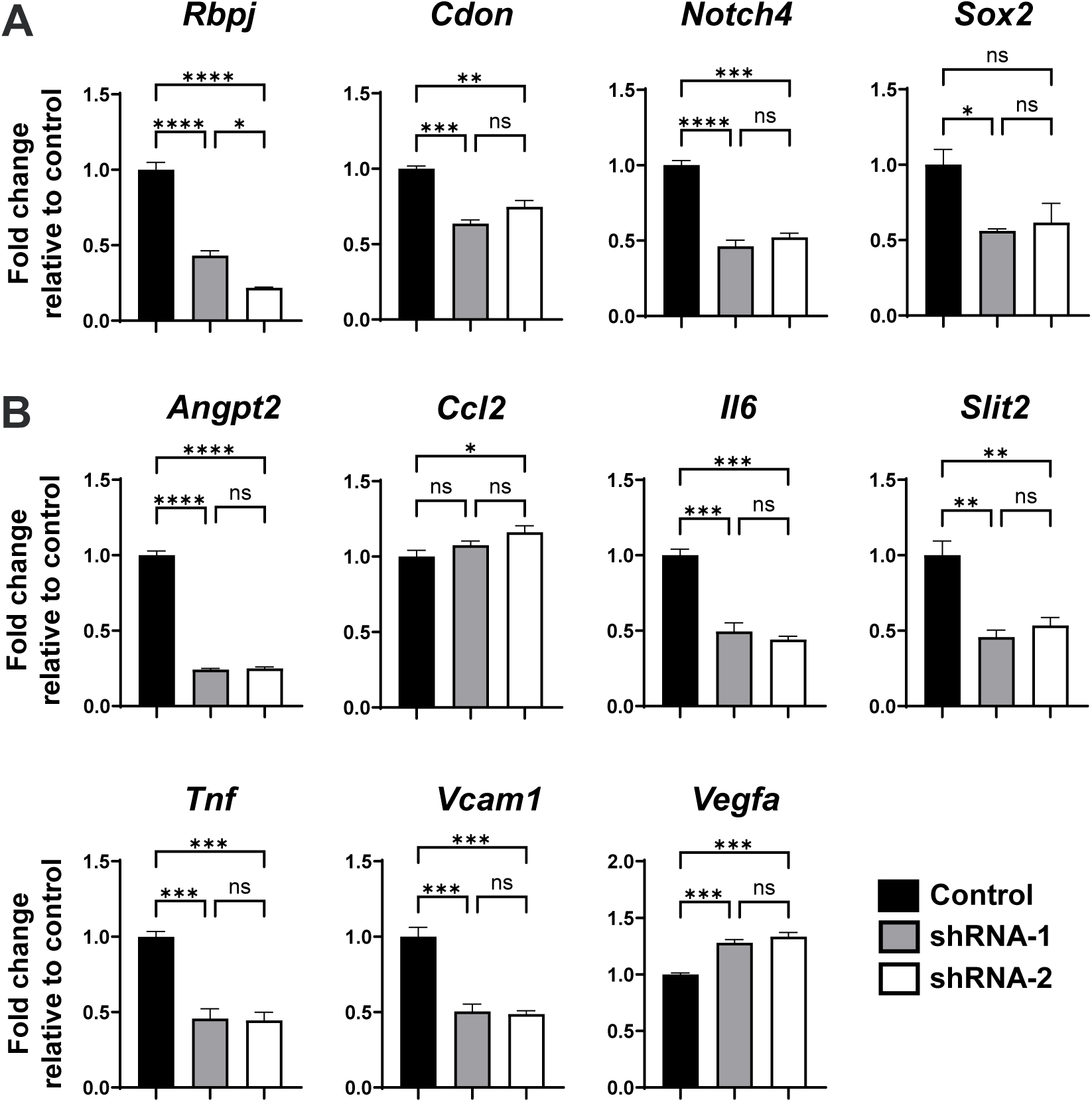
*Rbpj* knockdown downregulates expression of Notch/RBPJ target genes and markers of reactive cholangiocytes. qPCR analysis was performed using HPCs transduced with lentiviral vectors expressing two different shRNAs against *Rbpj* (shRNA-1 and shRNA-2) and a non-silencing control and then subjected to puromycin selection. (A) Downstream targets of Notch/RBPJ signaling. (B) Markers of reactive cholangiocytes. All data are expressed as mean ± SE; n = 3 independent experiments. * p < 0.05; ** p < 0.01; *** p < 0.001; **** p < 0.0001; ns, not significant. Abbreviations: HPC, hepatic progenitor cell; qPCR, quantitative polymerase chain reaction.

Since our data suggest that *Rbpj*-deletion in CK19+YFP+ HPCs affects CK19+YFP-cholangiocyte number (Figure 2), we hypothesized that modulation of CK19+YFP+ HPCs attenuates not only overall pathogenic progression but also activation of the cholangiocyte compartment in DDC-fed mice. To investigate this further, we chose vascular cell adhesion molecule 1 (VCAM1) for subsequent immunofluorescence analysis (Figure 7A) because of its roles in liver fibrosis and inflammation.[41] VCAM1-positive area and the number of CK19+VCAM1+ cells were induced upon DDC treatment in WT, and were significantly reduced in CKO (Figures 7B, C). We did not detect any CK19+ cells that express VCAM1 in chow-fed animals (Figure 7C). In DDC-fed mice, the percentage of VCAM1-expressing cells decreased in CKO compared to WT, not only in CK19+YFP+ HPCs but also in CK19+YFP-cholangiocytes (Figure 7D). Our data indicate that conditional *Rbpj* deletion specifically in HPCs not only inhibits reactive cholangiocyte marker expression in HPCs but also suppresses pathogenic activation of cholangiocytes that are not labeled by *Foxl1-Cre*.

**Figure 7.**
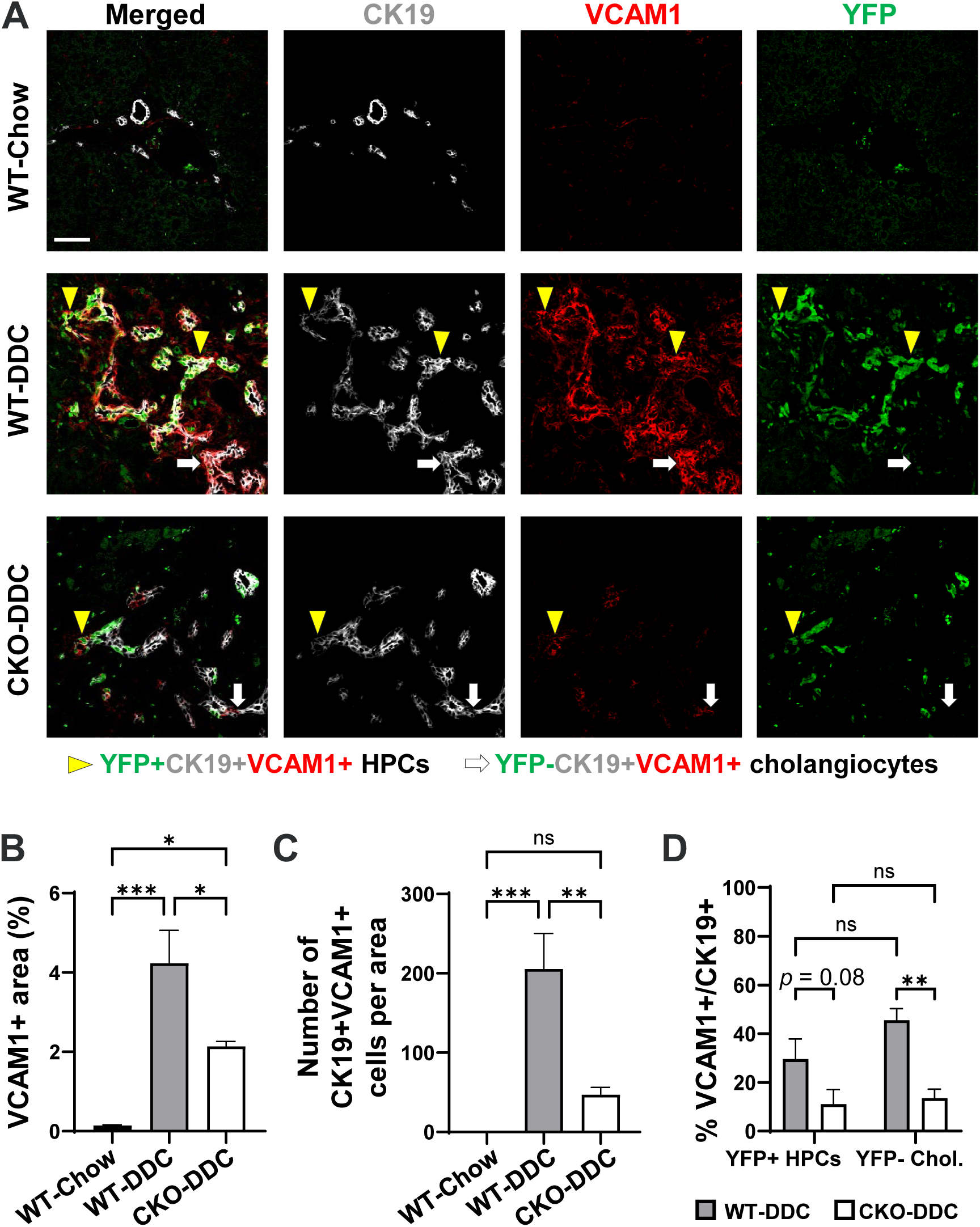
The percentage of VCAM1+ cells is reduced in both HPC and cholangiocyte populations in CKO-DDC compared to WT-DDC. (A) Confocal immunofluorescence analysis of portal areas. Yellow arrowheads: YFP+CK19+VCAM1+ HPCs. White arrows: YFP-CK19+VCAM1+ cholangiocytes. Scale bar, 50 μm. (B) Quantification of VCAM1+ portal areas. (C) Number of CK19+VCAM1+ cells per portal area. (D) Percentages of VCAM1+ cells in relation to the total number of CK19+YFP+ HPCs and CK19+YFP-cholangiocytes (Chol.) in the portal areas of WT-DDC and CKO-DDC livers. All data are expressed as mean ± SE; n = 5 mice per group. * p < 0.05; ** p < 0.01; *** p < 0.001; ns, not significant. Abbreviations: CK19, cytokeratin 19; CKO, conditional knockout; DDC, 3,5-diethoxycarbonyl-1,4-dihydrocollidine; HPC, hepatic progenitor cell; VCAM1, vascular cell adhesion molecule 1; WT, wild-type; YFP, yellow fluorescent protein.

In summary, our data suggest that conditional deletion of *Rbpj* specifically in *Foxl1-Cre*/YFP-labeled HPCs leads to reduced liver injury, vascular responses, and fibrosis in DDC-fed mice, and suppresses reactive phenotypes of YFP+ HPCs and YFP-cholangiocytes.

## DISCUSSION

This study aimed to determine the causal role of HPCs in pathogenic progression upon DDC treatment and the requirement of RBPJ in this process. The use of the *Foxl1-Cre* transgenic line allowed us to specifically delete *Rbpj* in HPCs. 21.7% of YFP-labeled HPCs in CKO-DDC maintained RBPJ expression. Because Cre efficiency is influenced by the position and distance between loxP sites, our data suggest that *Foxl1* promoter-driven Cre recombinase may induce YFP expression and delete *Rbpj* at different efficiencies.[42] Nevertheless, this level of gene ablation in HPCs was sufficient to induce phenotypic differences between WT-DDC and CKO-DDC. Our approach led to a significant finding that (1) HPCs are capable of remodeling the portal vascular and fibrotic microenvironment in response to chronic liver disease; (2) conditional modulation of HPCs was sufficient to inhibit overall pathogenic progression and to attenuate ductular reactions, liver injury, vascular expansion, and fibrosis in DDC-fed mice; and (3) RBPJ is required for the expression of reactive cholangiocyte markers in *Foxl1-Cre*/YFP-labeled HPCs and their ability to communicate with YFP-negative cholangiocytes.

Here, we use sNuc-seq as a proof-of-principle approach to demonstrate that conditional genetic manipulation of HPCs alters gene expression in multiple cell types. Analysis of each cluster at a higher resolution and experimental validation will be necessary for an accurate interpretation of the data. Our approach is limited, as sNuc-seq does not provide spatial information. A potential explanation for the marginal difference in the percentage of endothelial cells between WT-DDC and CKO-DDC is that sNuc-seq captures cells from all liver zones, whereas our immunofluorescence analysis focused specifically on portal areas. We chose sNuc-seq instead of single-cell RNA sequencing because hepatocyte populations in the DDC model were not represented efficiently using the latter method according to a study published by Wang and colleagues.[43] The hepatocyte representation was improved in our datasets, confirming the validity of our method, but Hepatocytes-1 cluster would still require experimental validation as it displays relatively weak enrichment of hepatocyte markers, likely due to the fact that the gene list used for annotation is based on uninjured liver.[32]

Another limitation of sNuc-seq analysis is that *Foxl1*-expressing HPCs were not detected in our sequencing data. To address this issue, we revisited our previously published microarray data[9] to demonstrate that HPCs are enriched for various markers of reactive cholangiocytes. Additionally, we treated an HPC line with anti-*Rbpj* shRNAs to demonstrate that HPC expression of many of these markers is dependent on *Rbpj*. Our data also indicate that HPC modulation impacts cholangiocyte number and VCAM1 expression, suggesting that attenuated activation of cholangiocytes could be one of the mechanisms explaining phenotypic differences between WT and CKO animals. The potential mechanisms behind this effect include: (1) attenuation of overall pathogenic progression in CKO mice leading to suppression of cholangiocyte activation and/or (2) paracrine regulation of cholangiocyte VCAM1 expression by inflammatory cytokines secreted by HPCs including tumor necrosis factor,[44] whose mRNA expression in HPCs was dependent on RBPJ (Figure 6). While it has been reported that Notch signaling regulates expression of some of reactive cholangiocyte markers such as interleukin 6 and VCAM1 in endothelial cells and macrophages,[45,46] our findings that *Rbpj* deletion in HPCs inhibits expression of reactive cholangiocyte markers in HPCs and also impacts cholangiocyte number and expression of VCAM1 are novel. The percentage of Ki67+ cells was high in WT HPCs compared to the CKO counterpart, but the difference did not reach significance in the cholangiocyte population. Possible explanations include that Ki67 only captures a snapshot of cells undergoing proliferation when tissues are harvested, or HPCs could affect cholangiocyte survival in addition to proliferation.

Immunostaining analysis indicated the presence of YFP-labeled hepatocytes in DDC-treated livers, and the fraction of mice with these labeled hepatocytes increased upon *Rbpj* deletion in HPCs (data not shown). However, whether lineage-labeled hepatocytes are detected or not did not correlate with any of the serum or immunofluorescence data analyzed in the current study, suggesting that the presence of labeled hepatocytes does not explain the phenotypic differences between WT and CKO mice. Rather, our data suggest that inhibition of overall pathogenic progression in CKO mice could explain attenuated vascular responses. Given that HPCs express many secretory factors involved in various pathogenic mechanisms, it is possible that HPCs also communicate with other cell types in addition to cholangiocytes, including hepatic stellate cells, hepatocytes, or immune cells to alter their pathogenic activation or recruitment, detailed mechanisms of which will be a subject of future research. While the mRNA levels of *Il6* and *Tnf* were decreased in the livers of CKO-DDC versus WT-DDC as well as in shRNA-treated HPCs versus control HPCs in vitro, the hepatic mRNA level of *Ccl2* did not correlate with in vitro data. This is likely because multiple cell types in addition to HPCs contribute to the total hepatic pool of secretory factors. Additionally, while RBPJ mediates Notch signaling when a ligand activates the pathway, it may act as a transcriptional repressor in a Notch-independent manner as well.[47] Therefore, de-repression of RBPJ may also be involved in phenotypes observed in CKO mice.

In conclusion, our study established the cause-and-effect relationship between HPCs and remodeling of the portal microenvironment, and demonstrated that RBPJ in HPCs is a suitable target for inhibiting not only ductular reactions and endothelial responses but also overall pathogenic progression. While our study is limited to a single disease model, it has been well-documented that cell-autonomous expression of Notch signaling components by biliary cells is implicated in the progression of several liver diseases, including primary sclerosing cholangitis, primary biliary cholangitis, polycystic liver disease, and cholangiocarcinoma.[48] Since the DDC diet induces both hepatocytic and cholestatic injuries, further investigation is warranted to determine whether the extent of HPC impact on pathogenic progression depends on the type of liver injury. The common occurrence of ductular reactions in many liver diseases[3] implies the potential for broad impact of our findings.

## ACKNOWLEDGEMENTS

We thank the Single Cell Genomics Facility [RRID# SCR_022653] and the Information Services for Research [RRID# SCR_022622] at Cincinnati Children’s Hospital Medical Center for sequencing and bioinformatic analysis services. We thank Dr. Tasuku Honjo for providing the *Rbpj^F/F^*mouse model used in this study.

## Author Contributions

Sanghoon Lee, Lu Ren, and Soona Shin conceptualized the work, designed experiments, interpreted the data, and wrote the initial draft. Sanghoon Lee, Lu Ren, Weiwei Li, and Ping Zhou conducted experiments. Stacey S. Huppert and Andrew Potter contributed to the optimization of experiments and design of the study. Sanghoon Lee, Lu Ren, Weiwei Li, and Aditi Paranjpe contributed to data acquisition, analysis, and visualization. All authors contributed to reviewing and editing the manuscript.

## Data availability

Raw data for single-nucleus RNA sequencing are in the process of being deposited in Gene Expression Omnibus (accession number to be added upon acceptance).

## Abbreviations

αSMA: alpha smooth muscle actin
ALP: alkaline phosphatase
AST: aspartate aminotransferase
CK19: cytokeratin 19
CKO: conditional knockout
DDC: 3,5-diethoxycarbonyl-1,4-dihydrocollidine
Foxl1: forkhead box L1
GFAP: glial fibrillary acidic protein
GSEA: gene set enrichment analysis
HPC: hepatic progenitor cell
HSC: hepatic stellate cell
LSEC: liver sinusoidal endothelial cell
NPC: non-parenchymal cell
qPCR: quantitative polymerase chain reaction
RBPJ: recombination signal binding protein for immunoglobulin kappa J region
sNuc-seq: single-nucleus RNA sequencing
VCAM1: vascular cell adhesion molecule 1
WT: wild-type
YFP: yellow fluorescent protein

