## Supplemental Information for "*Rbpj* deletion in hepatic progenitor cells attenuates endothelial responses and fibrosis in DDC-fed mice"

### This document includes:

**Supplemental Figure S1.** sNuc-seq reveals the impact of conditional HPC modulation on multiple cell types in the liver.

**Supplemental Figure S2.** Microarray analysis indicates enrichment of reactive cholangiocyte markers in HPCs compared to other cell types.

**Supplemental Materials and Methods**

### Supplemental_Tables.xlsx includes:

**Supplemental Table S1.** Primer sequences used for qPCR.

**Supplemental Table S2.** Cluster-specific markers.

**Supplemental Table S3.** List of overlapping markers.

**Supplemental Table S4.** Gene set enrichment analysis of markers for hepatocyte clusters.

**Supplemental Table S5.** Differentially expressed genes in each cluster.

**Supplemental Table S6.** Gene set enrichment analysis of differentially expressed genes in Cluster 0.

**Supplemental Table S7.** Gene set enrichment analysis of differentially expressed genes in Cluster 1.

**Supplemental Table S8.** Gene set enrichment analysis of differentially expressed genes in Cluster 2.

**Supplemental Table S9.** Gene set enrichment analysis of differentially expressed genes in Cluster 3.

**Supplemental Table S10.** Gene set enrichment analysis of differentially expressed genes in Cluster 4.

**Supplemental Table S11.** Gene set enrichment analysis of differentially expressed genes in Cluster 5.

**A**

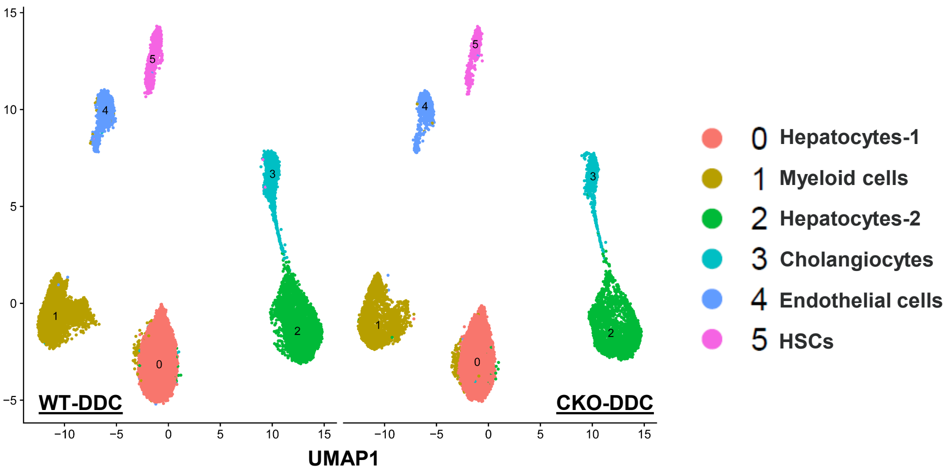


**B**

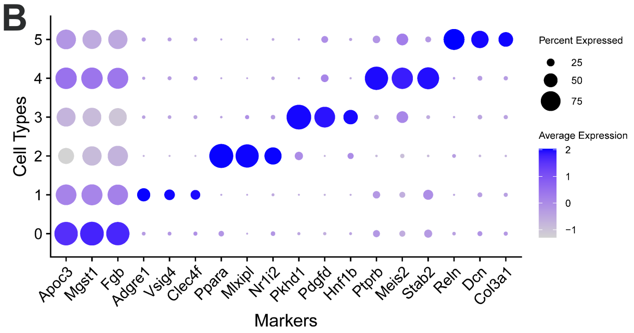


**C**

| Cluster | Population | Percentage WT-DDC (n = 2) | Percentage CKO-DDC (n = 2) |
|---|---|---|---|
| 0 | Hepatocytes-1 | 49.2 | 48.6 |
| 1 | Myeloid cells | 15.5 | 14.7 |
| 2 | Hepatocytes-2 | 16.4 | 23.2 |
| 3 | Cholangiocytes | 7.9 | 4.0 |
| 4 | Endothelial cells | 6.6 | 5.9 |
| 5 | HSCs | 4.4 | 3.6 |

**D**

**Cluster 3 Cholangiocytes**

| Name | Gene sets | NES | FDR q-val |
|---|---|---|---|
| BILANGES_SERUM_AND_RAPAMYCIN_SENSITIVE_GENES | M2 | -2.4070660 | 0.0000407 |
| BILANGES_RAPAMYCIN_SENSITIVE_VIA_TSC1_AND_TSC2 | M2 | -2.1040510 | 0.0090434 |
| REACTOME_RECOGNITION_OF_DNA_DAMAGE_BY_PCNA_ CONTAINING_REPLICATION_COMPLEX | M2 | -1.8863353 | 0.0954432 |
| REACTOME_SUMOYLATION_OF_DNA_REPLICATION_PROTEINS | M2 | -1.8173423 | 0.1617337 |
| GOCC_CENTRIOLE | M5 | -1.8696686 | 0.1898429 |
| GOMF_TRANSMEMBRANE_RECEPTOR_PROTEIN_KINASE_ACTIVITY | M5 | -1.8344204 | 0.2499141 |

**Cluster 4 Endothelial cells**

| Name | Gene sets | NES | FDR q-val |
|---|---|---|---|
| REACTOME_DNA_REPLICATION | M2 | -1.6639255 | 0.1770254 |
| REACTOME_KEAP1_NFE2L2_PATHWAY | M2 | -1.6531448 | 0.1737312 |
| HEVNER_CORTEX_VASCULAR_ENDOTHELIAL_CELLS | M2 | -1.6351291 | 0.1994533 |
| REACTOME_SIGNALING_BY_HEDGEHOG | M2 | -1.6311255 | 0.2001589 |
| REACTOME_SIGNALING_BY_VEGF | M2 | -1.6289442 | 0.1970500 |
| GOBP_VASCULAR_ENDOTHELIAL_GROWTH_FACTOR_SIGNALING_ PATHWAY | M5 | -1.8825284 | 0.0328418 |
| GOBP_CELLULAR_RESPONSE_TO_VASCULAR_ENDOTHELIAL_ GROWTH_FACTOR_STIMULUS | M5 | -1.7797999 | 0.1316190 |

**Cluster 5 HSCs**

| Name | Gene sets | NES | FDR q-val |
|---|---|---|---|
| REACTOME_COLLAGEN_CHAIN_TRIMERIZATION | M2 | -1.9520935 | 0.0624917 |
| REACTOME_O_LINKED_GLYCOSYLATION | M2 | -1.8382125 | 0.1170048 |
| NABA_COLLAGENS | M2 | -1.7581775 | 0.1820221 |
| NABA_ECM_AFFILIATED | M2 | -1.6909060 | 0.2496825 |
| GOBP_INTEGRIN_MEDIATED_SIGNALING_PATHWAY | M5 | -1.9220525 | 0.1653669 |
| GOBP_EXTRACELLULAR_STRUCTURE_ORGANIZATION | M5 | -1.9050715 | 0.1822949 |

**Supplemental Figure S1. sNuc-seq reveals the impact of conditional HPC modulation on multiple cell types in the liver.** Livers from WT-DDC (n = 2) and CKO-DDC (n = 2) were subjected to sNuc-seq analysis. (A) UMAP plots showing 6 clusters. (B) Representative markers of each cluster. (C) The percentage of nuclei for each cluster. The average value for 2 samples in each genotype is shown. (D) GSEA was performed using genes downregulated in CKO-DDC compared to WT-DDC in each cluster and selected enriched terms are shown. An FDR q-value of <0.25 was considered statistically significant in this study. Abbreviations: CKO, conditional knockout; DDC, 3,5-diethoxycarbonyl-1,4-dihydrocollidine; FDR, false discovery rate; GSEA, Gene set enrichment analysis; HSC, hepatic stellate cell; NES, normalized enrichment score; sNuc-seq, single-nucleus RNA sequencing; UMAP, uniform manifold approximation and projection; WT, wild-type.

**A**

| | Cholangiocytes-Chow /HPCs-DDC | | Hepatocytes-Chow /HPCs-DDC | |
|---|---|---|---|---|
| **Gene** | **Fold change** | **FDR** | **Fold change** | **FDR** |
| *Angpt1* | *-17.54* | 2.27 | *-41.67* | 0 |
| *Angpt2* | *-71.43* | 0.93 | *-17.24* | 0 |
| *Ccl2* | *-125.00* | 0.96 | *-Inf* | 0 |
| *Ctgf* | -1.11 | 53.42 | *-Inf* | 0 |
| *Cxcl1* | *-4.63* | 3.63 | *-2.09* | 1.71 |
| *Cxcl16* | 4.23 | 3.63 | *-14.29* | 0 |
| *Il1b* | -2.62 | 100.00 | 1.39 | 12.77 |
| *Il4* | *-2.53* | 10.55 | *-1.42* | 4.22 |
| *Il6* | -1.41 | 100.00 | 2.93 | 1.17 |
| *Il7* | 1.23 | 55.75 | *-15.63* | 0 |
| *Il10* | -1.92 | 40.59 | 1.06 | 21.49 |
| *Il18* | 7.21 | 4.52 | 10.31 | 0 |
| *Itgb6* | 5.43 | 2.48 | *-23.81* | 0 |
| *Pdgfb* | *-4.00* | 3.98 | *-58.82* | 0 |
| *Serpine1* | -1.49 | 44.86 | *-250.00* | 0 |
| *Spp1* | *-24.39* | 0.66 | *-1000.00* | 0 |
| *Tgfb1* | *-1.59* | 10.55 | *-13.51* | 0 |
| *Tgfb2* | *-6.58* | 6.29 | *-166.67* | 0 |
| *Tnf* | 15.09 | 2.80 | *-2.53* | 0.81 |
| *Vcam1* | *-8.55* | 2.27 | *-333.33* | 0 |
| *Vegfa* | 5.83 | 0.79 | 6.47 | 0 |

**B**

| | NPCs-DDC /HPCs-DDC | |
|---|---|---|
| **Gene** | **Fold change** | **FDR** |
| *Angpt1* | *-4.61* | 17.18 |
| *Angpt2* | -1.34 | 49.46 |
| *Ccl2* | *-4.46* | 5.76 |
| *Ctgf* | 1.07 | 61.19 |
| *Cxcl1* | *-2.62* | 7.42 |
| *Cxcl16* | *-1.91* | 8.02 |
| *Il1b* | 2.92 | 23.84 |
| *Il4* | *-4.79* | 11.84 |
| *Il6* | 11.33 | 21.55 |
| *Il7* | *-5.78* | 10.32 |
| *Il10* | *-4.95* | 5.44 |
| *Il18* | *-2.39* | 12.88 |
| *Itgb6* | *-8.33* | 7.42 |
| *Pdgfb* | *-2.77* | 8.92 |
| *Serpine1* | 4.67 | 26.32 |
| *Spp1* | *-3.34* | 6.54 |
| *Tgfb1* | 1.81 | 44.15 |
| *Tgfb2* | *-3.16* | 8.02 |
| *Tnf* | *-5.99* | 8.02 |
| *Vcam1* | *-2.31* | 12.88 |
| *Vegfa* | *-4.17* | 7.42 |

**Supplemental Figure S2. Microarray analysis indicates enrichment of reactive cholangiocyte markers in HPCs compared to other cell types.** Expression profiling data from GSE28892 are shown. All cells were isolated from wild-type male mice. (A) Primary cholangiocytes and primary hepatocytes from chow-fed mice were compared to HPCs from mice fed the DDC diet for 2 weeks. (B) NPCs collected after exclusion of HPCs and hepatocytes were compared to HPCs from the same animals fed with the DDC diet for 2 weeks. All data are expressed as expression levels in relation to HPCs. Statistically significant upregulation in HPCs compared to other populations is marked by the use of underlined and italicized numbers. Genes whose expression was undetectable in hepatocytes are marked '-Inf'. FDR < 20% was considered statistically significant in this study. 3-4 replicates per each condition were analyzed except for primary cholangiocytes with 2 replicates. Abbreviations: DDC, 3,5-diethoxycarbonyl-1,4-dihydrocollidine; FDR, false discovery rate; HPC, hepatic progenitor cell; NPC, non-parenchymal cell.

### Supplemental Materials and Methods

#### *Single-nucleus RNA sequencing*

Nuclei were prepared from fresh-frozen WT-DDC (n = 2) and CKO-DDC (n = 2) mouse livers using the protocol published by Humpherys and Kirita[1,2] with a modification: Addition of 50 μL of 4% paraformaldehyde to the homogenate after passing through a cell strainer.[3] The single-nucleus RNA sequencing (sNuc-seq) assay was performed according to the manufacturer's instructions (Chromium Next GEM Single Cell 3' Reagent Kits v3.1 (Dual Index), 10x Genomics). Briefly, nuclei were resuspended in the master mix and loaded together with partitioning oil and gel beads into the chip to generate a gel bead-in-emulsion. The poly-A RNA from the cell lysate contained in every gel bead-in-emulsion was reverse transcribed to cDNA, adding an Illumina R1 primer sequence, unique molecular identifiers (UMIs), and the 10x Barcode. The barcoded cDNA was then cleaned up with Silane DynaBeads and amplified for 18 cycles. Full-length, barcoded cDNA was then enzymatically fragmented, sized-selected using SPRIselect reagent, adapter-ligated, and amplified for library construction. During library construction, Illumina R2 primer sequence, paired-end constructs with P5 and P7 sequences, and sample indexes were added. Libraries were pooled and sequenced on the NovaSeq 6000 using S4 flow cells with the following sequencing parameters: R1: 28 cycles, i7: 10 cycles, i5: 10 cycles, R2: 90 cycles.

#### *Bioinformatic analysis of sNuc-seq data*

Raw base call files were de-multiplexed with Cell Ranger v5.0.1 mkfastq.[4] Reads were aligned to mouse reference genome mm10 and gene expression was quantified using Cell Ranger count. Ambient RNA fraction was estimated and removed using SoupX v1.5.2[5] with the autoEstCont and adjustCounts functions. Further data analysis was carried out with Seurat v4.9.9 in R 4.1.1.[6-8] Cells with more than 5% mitochondrial gene expression, less than 500 total expressed genes, less 1000 UMIs were excluded from the analysis. UMI counts were normalized with the SCTransform Seurat function. The four samples were integrated together using FindIntegrationAnchors and IntegrateData functions from Seurat. Principal component analysis (PCA) was performed using RunPCA Seurat function (the number of principal components = 40). Uniform manifold approximation and projection (UMAP) was run using the top 40 PCA vectors as input to RunUMAP Seurat function. Cells were clustered using the FindNeighbors function (top 40 PCA vectors) and FindClusters function (with a range of possible resolutions between 0 to 1). Marker genes for each cluster were identified using FindConservedMarkers Seurat function with default parameters. A resolution value of 0.1 was ultimately selected, as distinct gene expression profiles were present in each cluster, while higher resolutions produced adjacent clusters with a high degree of overlapping marker genes. Differentially expressed genes (DEGs) for each cluster were identified using the Wilcoxon rank sum test with FindMarkers function. The gene should be expressed in a minimum 10% of the cells on either side. The list of DEGs and log2 fold changes were further used for gene set enrichment analysis (GSEA). GSEAPreranked tool within GSEA v3.0.0[9,10] was applied with M2 and M5 gene sets from the Molecular Signatures Database (MSigDB).[11,12]
